# Quorum quenching activity of the PGPR *Bacillus subtilis* UD1022 alters nodulation efficiency of *Sinorhizobium meliloti* on *Medicago truncatula*

**DOI:** 10.1101/2020.08.26.268953

**Authors:** Amanda Rosier, Pascale B. Beauregard, Harsh P. Bais

## Abstract

Plant growth promoting rhizobacteria (PGPR) have enormous potential for solving some of the myriad challenges facing our global agricultural system. Intense research efforts are rapidly moving the field forward and illuminating the wide diversity of bacteria and their plant beneficial activities. In the development of better crop solutions using these PGPR, producers are including multiple different species of PGPR in their formulations in a ‘consortia’ approach. While the intention is to emulate more natural rhizomicrobiome systems, the aspect of bacterial interactions has not been properly regarded. By using a tri-trophic model of *Medicago truncatula* A17 Jemalong, its nitrogen (N)-fixing symbiont *Sinorhizobium meliloti* Rm8530 and the PGPR *Bacillus subtilis* UD1022, we demonstrate indirect influences between the bacteria affecting their plant growth promoting activities. Co-cultures of UD1022 with Rm8530 significantly reduced Rm8530 biofilm formation and downregulated quorum sensing (QS) genes responsible for symbiotically active biofilm production. This work also identifies the presence and activity of a quorum quenching lactonase in UD1022 and proposes this as the mechanism for non-synergistic activity of this model ‘consortium’. These interspecies interactions may be common in the rhizosphere and are critical to understand as we seek to develop new sustainable solutions in agriculture.

## Introduction

Global agriculture is facing extraordinary challenges driven by factors associated with current intensive management practices. These problems include and are not limited to the unsustainable use of synthetic and extracted fertilizers, destruction of ecosystems for more arable land, contributions to CO_2_ and methane emissions and the consequences of global climate change (Tilman et al., 2011). Many of these issues arise, in part, due to the demands of a growing population, with the 2019 UN *Highlight* reporting that we will have between 9.4 and 10.1 billion people on earth by 2050 (UN, 2019). Though crop productivity and yields are greater than any time in history, they are still not projected to meet the nutritional needs of this many people (Tilman et al., 2011).

An estimated 30-50% of current yields are estimated to be due to the use of synthetic fertilizer including N (Stewart et al., 2005). The sharp growth in the numbers of people who could be supported per hectare of arable land rose from 1.9 in 1908 to 4.3 in 2008 and is attributed primarily to N sourced from the Haber-Bosch process (Erisman et al., 2008). Haber-Bosch N production requires high amounts of energy to break the triple bond between N_2_ to form plant available NH_3_ (ammonia). About 1% of global fossil energy is consumed by Haber-Bosch to manufacture synthetic N (Blaser et al., 2016) and 80% of the N produced goes toward agriculture fertilizers (Galloway et al., 2008). Shockingly, of the N created, only 2-10% reaches our table (Fields, 2004). This inefficiency of N use in agriculture stems from its rapid loss to the environment via multiple avenues before it is taken up by crops (Robertson and Vitousek, 2009). Excess N poses a multitude of risks to environmental and human health as detailed in several reviews and analysis (Zhang et al., 2015; Houlton et al., 2019; Liu et al., 2020).

Currently, a multitude of efforts and approaches are being studied to address the greatest problems inherent to large-scale agriculture. Solving these problems will likely require a combination of many strategies including crop breeding, plant genetic biotechnology, sustainable field management, alternative crops, and soil health improvements (Gouda et al., 2018). One solution which shows great promise in improving soil and plant health is to harness the natural plant microbiome found inhabiting the ecosystem on and around plant roots, termed the ‘rhizosphere’ (Adesemoye et al., 2009; Gamalero and Glick, 2011). The organisms isolated from the rhizosphere which improve plant health and resilience directly or indirectly are known as plant growth-promoting rhizobacteria (PGPR) (Kloepper et al., 1980). PGPRs have a variety of lifestyles including as endosymbionts and existing in microcolonies and biofilms on the root surface (Gray and Smith, 2005; Beauregard, 2015). They provide direct and indirect benefits to the plant through niche competition with other microbes, production of antibiotics to inhibit pathogens, mineralization of organic macronutrients, chelation of micronutrients, and stimulation of the plant’s own immune response systems to defend it from invading pathogens (Gamalero and Glick, 2011; Mendes et al., 2011).

The classic example of plant-microbe partnerships is the mutualism between symbiotic N-fixing bacteria in the family *Rhizobia* and their specific legume plant hosts. In a process referred to as ‘biological nitrogen fixation’ (BNF), *Rhizobia* fix atmospheric N for the plant in exchange for carbon-rich photosynthates within specialized structures formed on the plant root called nodules (Oldroyd, 2013). Legume crops are important in sustainable agriculture for their multifaceted benefits to ecology and human health (Stagnari et al., 2017). Peoples et al. (2009) estimates that 30-40 kg of N is fixed for every ton of crop legume dry matter and Herridge et. al. (2008) approximated that N-fixation by crop and forage legumes through symbiosis globally is roughly 50 Tg per year. This ability of *Rhizobia* to fix nitrogen allows for a de-coupling from the dependence on synthetic nitrogen. As such, it is the focus of intense efforts to engineer non-legumes to either be compatible with N-fixing bacteria or to incorporate the molecular N-fixing pathway into cereal crops (Rogers and Oldroyd, 2014; Pankievicz et al., 2019).

Products utilizing live PGPR or microbially derived metabolites for crop application have become a growing proportion of agrochemical company research efforts. Broadly termed ‘biologicals’, these products can be subcategorized as biopesticides and biostimulants for crop protection against pests and pathogens and as biofertilizers consisting of bacteria which solubilize phosphorous, fix N_2_ or otherwise make essential nutrients more available for the plant (Timmusk et al., 2017; Marrone, 2019). Global markets for these products is growing, with estimates of the current value of microbial soil inoculants at around USD 396 million and growing at an annual rate of 9.5% and biofertilizers expected to reach USD 2653.48 million by 2023 (Arora et al., 2020). These products are proving to have so much potential that new market formulations are focusing on using multiple different species of live bacteria in ‘consortia’ (Marrone, 2019). The rationale behind this innovation derives from the knowledge that natural rhizomicrobiomes are occupied by millions of different species of bacteria working in conjunction with one another (Schlatter et al., 2015); restoration of these microbial ecosystems may provide more robust benefits to the plant (Gouda et al., 2018; Sergaki et al., 2018).

Synergistic plant growth promotion through the presence of multiple PGPR species has been observed in certain plant – bacteria – bacteria systems (Schwartz et al., 2013; Morel et al., 2015; Berendsen et al., 2018), but have also failed in others (Felici et al., 2008; Kang et al., 2014; Maymon et al., 2015). Screening for beneficial plant outcomes in the search for compatible PGPR interspecies interactions generally ignores underlying mechanisms responsible for enhanced plant responses. An important aspect of identifying suitable PGPR consortia will be to understand the multitude of plant beneficial activities that may be altered when the organisms coexist in what is now more commonly being described as the plant holobiont (Zilber-Rosenberg and Rosenberg, 2008). The dearth of identified PGPR interspecies interactive mechanisms is a significant gap in our knowledge but poses an opportunity to make empirical selections of appropriate PGPRs as we continue to expand our understanding of their plant beneficial activities.

To investigate meaningful plant – PGPR mechanisms, we designed a simplified tri-trophic legume – symbiont – PGPR system consisting *of Medicago truncatula* A17 Jemalong, its symbiotic mutualist *Sinorhizobium meliloti* strain Rm8530 and the PGPR *Bacillus subtilis* strain UD1022 (Rosier, 2016). Similar combinations of legumes, symbionts and PGPRs have been evaluated, however the experimental designs typically only report plant growth parameters and bacterial metabolite production without identifying or implicating specific interacting mechanisms between the species involved. The organisms in this model were specifically selected to be representative due to their comprehensively described genetics and lifestyles.

The genome of *M. truncatula* has been completely sequenced,is well annotated, and extensive mutant collections are available (Young et al., 2011; Krishnakumar et al., 2015). *M. truncatula* is also more genetically tractable than its relative *M. sativa* (alfalfa) but is similar enough to allow findings to be translated to the valuable perennial forage crop (Young and Udvardi, 2009). Alfalfa production is important as an N-fixing high protein forage for meat and dairy industries (Li and Brummer, 2012), sustainable crop rotations (Kulkarni et al., 2018), and stabilizing soil organic carbon (Li et al., 2019). In the United States, alfalfa was the third most valuable field crop with hay valued at USD 10.9 billion in 2019 (https://quickstats.nass.usda.gov, 2020).

Symbiosis between *M. truncatula* and *S. meliloti* is instigated through the exudation of the specific signaling plant flavonoid luteolin which acts as a chemoattractant (Hassan and Mathesius, 2012). More importantly, luteolin induces the transcription and expression of *S. meliloti nod* genes, producing lipo-chitooligosaccharide signals called Nod factors (NF) (Kondorosi et al., 1989). The symbiotic interspecies molecular signaling program is exquisitely complex and very well described (Oldroyd et al., 2011; Gourion et al., 2015; Zipfel and Oldroyd, 2017), including identification of the *M. truncatula* root hair localized LysM receptor-like kinases NFP and LYK3 recognition of *S. meliloti* specific NFs (Amor et al., 2003; Smit et al., 2007), subsequent infection thread development and commencement of nodule organogenesis (Limpens et al., 2003; Gage, 2004). *S. meliloti* then divide, proliferate and express N_2_-fixing nitrogenase enzyme within these plant-derived nodules (Oldroyd, 2013).

Importantly, certain *S. meliloti* exopolysaccharides (EPS) are known to be required for the successful initiation of nodules (Marketon et al., 2003). These EPS include succinoglycan and high and low molecular weight molecules of galactoglucan (EPS II) present in *S. melilotii* biofilms (Rinaudi and Gonzalez, 2009). The EPS II fraction of these biofilms are described as ‘symbiotically active’ as EPS II defective mutants are ineffective at forming WT pink nodules (González et al., 1996). Further analysis shows that EPS II production is dependent on the quorum sensing (QS) regulatory network including *expR* (Pellock et al., 2002) and *sinI* genes (Marketon et al., 2003). The *S. meliloti* ExpR/SinI QS system relies on SinI synthase-produced long chain N-acyl homoserine lactone (AHL) signal molecules (Marketon et al., 2002; Gao et al., 2005). AHL bound ExpR transcriptional response controls the expression of up to 500 different QS controlled genes (Gurich and González, 2009). The common lab model strain of *S. meliloti* Rm1021 has an insertion element in its ExpR gene. Since a complete QS biofilm system is critical for the successful invasion of nodules, the ExpR+ *S. meliloti* strain Rm8530 (Glazebrook and Walker, 1989; Marketon et al., 2003) was chosen for our tri-trophic model system.

Lastly, the University of Delaware patented *B. subtilis* strain UD1022 (Bais et al., 2015; Bishnoi et al., 2015) is incorporated into our tri-trophic model as an ideal generalist PGPR. A broad diversity of *Bacillus* species are known to have equally expansive and robust PGP activities. These activities include phytohormone production, nutrient solubilization and a multitude of plant protective features such as production of antifungal secondary metabolites and volatile organic compounds (VOCs) (Aloo et al., 2019). Further, some of these compounds, such as the cyclic lipopeptide (CLP) surfactin, play a dual role by also activating the plants own defense pathway known as induced systemic resistance (ISR) (Cawoy et al., 2014; Thomashow et al., 2019). As well as producing a rich variety of secondary metabolites and lytic enzymes, several *Bacillus* spp are also known to express quorum quenching (QQ) enzymes, which disrupt QS signaling of other bacteria, including pathogens (Dong et al., 2001; d’Angelo-Picard et al., 2005; Ryan et al., 2009). Details of *Bacillus* PGP activities from the lab to bioinoculants can be found in the following comprehensive reviews (Kumar et al., 2011; Chowdhury et al., 2015; Wu et al., 2015).

Various co-inoculation combinations of *Bacillus* species with legumes and their symbionts have shown positive growth and nodulation outcomes. Most prominently, nodulation was greatly enhanced in soybean (Glycine max L. Merr) systems when *Bradyrhizobium japonicum* was co-inoculated with *B. cereus* UW85 (Halverson and Handelsman, 1991), *B. thuringiensis* NEB17 (Bai et al., 2003) or with *B. amyloliquefaciens* strain LL2012 (Masciarelli et al., 2014). *B. subtilis* specific co-inoculations have been successful when used with *Rhizobium leguminosarum* bv. *viciae* 128C53 (Rlv) onto *Pisum sativum* L. (pea) (Schwartz et al., 2013) and when applied along with *Bradyrhizobium japonicum* on soybean (Bai et al., 2003). No direct mechanisms of interaction were tested in these studies; Schwartz et al. (2013) surveyed *B. simplex* 30N-5 and found possible fungal antagonism, phosphate solubilization, siderophore and auxin activities. In a follow-up study to attempt to expand their work, Maymon et al. (2015) applied *B. simplex* 30N-5 to an *M. truncatula* – *S. meliloti* 1021 system and found no significant plant growth improvement.

The primary goal of this work is to identify specific interspecies interactions between PGPR’s that influence their plant beneficial activities in the legume-rhizobium symbiosis. Through using the tri-trophic model as a platform for testing phenotypic outcomes of the ‘consortium’, more fundamental questions regarding the interspecies interactions can be developed. Specifically, does the PGPR and legume symbiont consortia act synergistically to increase *M. truncatula* plant growth, how do the different PGPR species directly or indirectly interact with one another, and do those interactions influence their ability to interact with and confer benefits to the plant? We employed phenotypic and molecular assays to evaluate the bacteria – bacteria interactions *in vitro* and *in vivo* on the plant root.

## Materials and Methods

### Bacterial Growth

Primary cultures of all bacteria strains were grown and maintained on TYC media (TY media (Beringer, 1974) liquid or agar supplemented with 1 mM CaCl_2_) with appropriate antibiotics. Subcultures of *Sinorhizobium meliloti* strain Rm8530 and *Bacillus subtilis* strain UD1022 (hereafter ‘UD1022) prepared for biofilm treatments were sub-cultured into minimal glutamate mannitol (MGM), low phosphate (0.1 mM), as described in (Marketon and Gonzalez, 2002). Both UD1022 and Rm8530 strains were grown at 30° C for all experiments. AT medium for culturing pre-induced *A. tumefaciens* KYC55 was prepared as described in Joelsson and Zhu, 2005.

### Plant Growth and Co-Inoculation

Seeds of *Medicago truncatula* A17 cv Jemalong were acid scarified for 6 minutes and sterilized with 3% bleach for 3 minutes. Seeds were imbibed in sterile water at 4° C overnight, rinsed and placed in sterile petri dish and germinated covered overnight at room temperature (Garcia et al., 2006). Germinated seeds were placed in sterile Magenta^®^ (Magenta Corp.) jars with Lullien’s solution (Lullien et.al., 1987), sealed with 3M^™^ MicroPore^™^ surgical tape and grown in a controlled environmental chamber at 55% relative humidity and a 14-h, 22° C day/10 h, 18°C night cycle. After 6 days of growth, plants were inoculated with bacteria treatments, with 10 plants per treatment. Rm8530 was grown to OD_600_ = 0.8 and UD1022 at OD_600_ = 1.0. Bacteria were spun down, washed 3 times in sterile H_2_O and re-suspended with 0.5X Lullein’s solution with Rm8530 final OD_600_ = of 0.02 and UD1022 OD_600_ = 0.01 (in Magenta jar). Plants were harvested 7 weeks after inoculation.

### Cross-streak for Growth Inhibition Analysis

Rm8530 bacteria were grown to OD_600_ = 0.8 and UD1022 OD_600_ = 1.5. Both cultures were diluted to OD_600_ = 0.5 with sterile H_2_O. Bacteria were streaked on TYC agar plates using a sterile loop in a cross pattern.

### Biofilm Assays

#### Preparation of Cell Free Supernatant (CFS) Derived from UD1022 for Biofilm Assays

UD1022 was inoculated from a single plate colony into 5 mL TYC and grown overnight (16 hr), then diluted 1:50 in 50 mL MGM in sterile 150 mL flask and grown shaking for 8 hours to an OD_600_ = 0.8 - 1.0. Cultures were centrifuged 10 minutes, 4° C at 4,000 RPM. Culture supernatant was filter-sterilized with 0.22 µm membrane (Steriflip^®^, EMD Millipore) under gentle vacuum. Supernatant was centrifuged and filter sterilized once more. A sub-fraction was heat treated in water bath overnight at 65° C.

#### Preparation of Biofilm Treatments

Biofilm assays were based on methods found in (O’Toole et al., 1999; Rinaudi and Gonzalez, 2009). Rm8530 was grown 48 hours in TYC to OD_600_ = 1.5-2.0, then cells were ‘pre-conditioned’ by diluting 1:100 to MGM media and grown shaking 48 hours to OD_600_ = 0.8. Stocks of treatments were made by centrifuging and re-suspending cell pellets with fresh MGM, or UD1022 CFS, UD1022 ‘heat treated’ CFS to a total of 5% by volume in MGM. 100 µL of these treatment stocks were then aliquoted to 96 well plates with 8 replicate wells per treatment. Plates were sealed with Parafilm^®^ (Bemis Company, Inc.) and placed in shaker at 30° C and measured at 24, 48 and 72 hours.

Experiment was repeated 3 separate times.

Plates were then emptied and gently rinsed 3 times with sterile water, dried, and stained 20 minutes with 150 µL of 0.1% crystal violet. Plates were emptied, rinsed gently 3 times with sterile water. Crystal violet (CV) was solubilized with modified biofilm dissolving solution (MBDS) (Tram et al., 2013). OD_570_ of CV was then measured using Wallac 1420 Plate Reader (PerkinElmer Life and Analytical Science, Wallac Oy, P.O. Box 10, FIN-20101 Tuku, Finland).

#### Gene Expression Reporter Assays

Reporter lines for Rm8530 were provided by Dr. Max Teplitski of the University of Florida. All cultures grown in liquid TYC broth shaking at 225 RPM at 30° C. Bacteria primary cultures were grown with appropriate antibiotics 48 hours to OD_600_ = 2.0-3.0. Cells were further prepared as described in ‘Biofilm Assays’ section and 29 replicate wells were included per treatment. Every 24 hours total well fluorescence and cell growth were measured using Wallac 1420 Plate Reader (PerkinElmer Life and Analytical Science, Wallac Oy, P.O. Box 10, FIN-20101 Tuku, Finland). Data reported as fluorescence counts/OD_570_ (Gao et al., 2012). After the 72-hour measurement, 96 well plates were processed as described in ‘Biofilm Assays’ above to assess qualitative biofilm formation. Gene reporter assays were repeated three times.

### Statistical Analysis

For plant growth biological data and biofilm analysis, data normality and homogeneity were reviewed prior to analysis of variance (ANOVA). No data transformations were required. One-way ANOVA was used to test for differences between treatments. When F-ratios were significant (p < 0.05), treatment means were compared via Tukey Kramer HSD using SAS-JMP (Cary, NC, USA).

For gene expression reporter results analysis, ANOVA was used to test for treatment differences. Where F-ratios were significant (P < 0:05), treatment means were compared via Tukey–Kramer test (JMP, SAS Institute). Non-parametric analyses (Kruskal–Wallis test) were utilized if data failed to meet parametric assumptions. Where H-values (Kruskal–Wallis test statistic) were significant (P < 0:05), treatment means were compared via Kruskal–Wallis multiple comparison Z-value test using NCSS software (Hintze, 2000).

### Gene Expression Analysis Using Semi-Quantitative Reverse Transcription PCR (qRT-PCR) Primer Design for qRT-PCR

Gene sequences were derived from GenBank; *S. meliloti* 1021 sequences from genome (accession: AL591688.1), and mega-plasmids pSymA (accession: AE006469.1). The *sinI* primer pair from (Gurich and González, 2009) and the *rpo*E1 primer pair from (Trabelsi et al., 2009). Primers from this work were designed using GenScript Real-time PCR (TaqMan) Primer Design (https://www.genscript.com/ssl-bin/app/primer). Amplicon size was restricted to 150 bp or less. All primer sequences (Table 2) were cross-checked on all strain sequences to ensure species specificity.

**TABLE 1.**
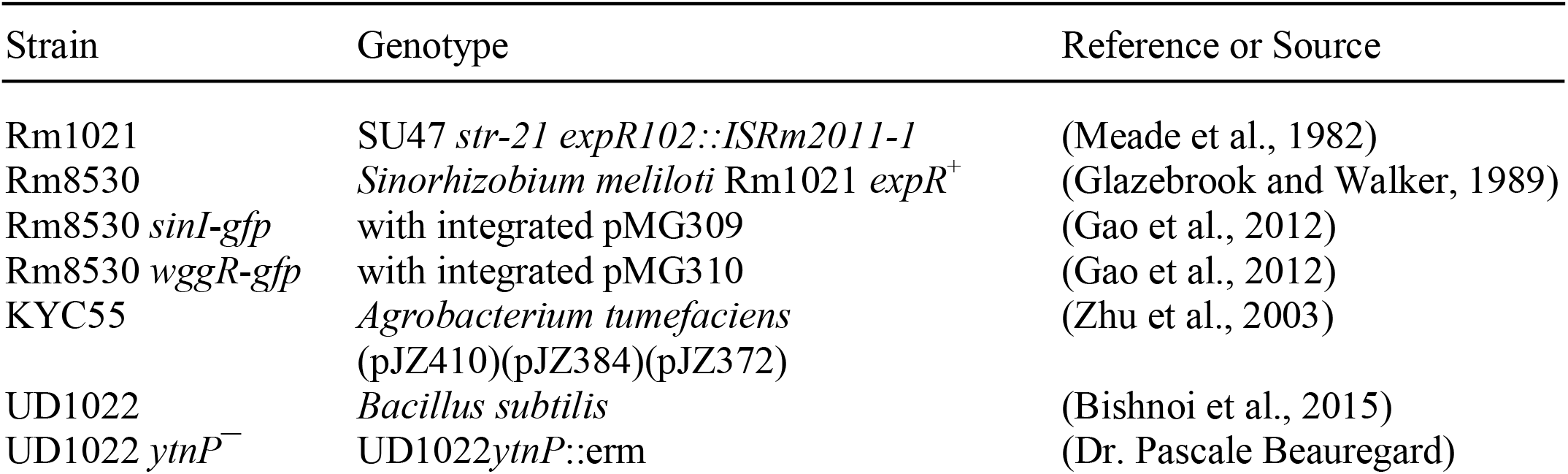
Bacterial strains used in this study.

| Strain | Genotype | Reference or Source |
| --- | --- | --- |
| Rm1021 | SU47 *str-21 expR102::ISRm2011-1* | (Meade et al., 1982) |
| Rm8530 | *Sinorhizobium meliloti* Rm1021 *expR*<sup>+</sup> | (Glazebrook and Walker, 1989) |
| Rm8530 *sinI-gfp* | with integrated pMG309 | (Gao et al., 2012) |
| Rm8530 *wggR-gfp* | with integrated pMG310 | (Gao et al., 2012) |
| KYC55 | *Agrobacterium tumefaciens*<br>(pJZ410)(pJZ384)(pJZ372) | (Zhu et al., 2003) |
| UD1022 | *Bacillus subtilis* | (Bishnoi et al., 2015) |
| UD1022 *ytnP*<sup>-</sup> | UD1022*ytnP::erm* | (Dr. Pascale Beauregard) |

**TABLE 2.**
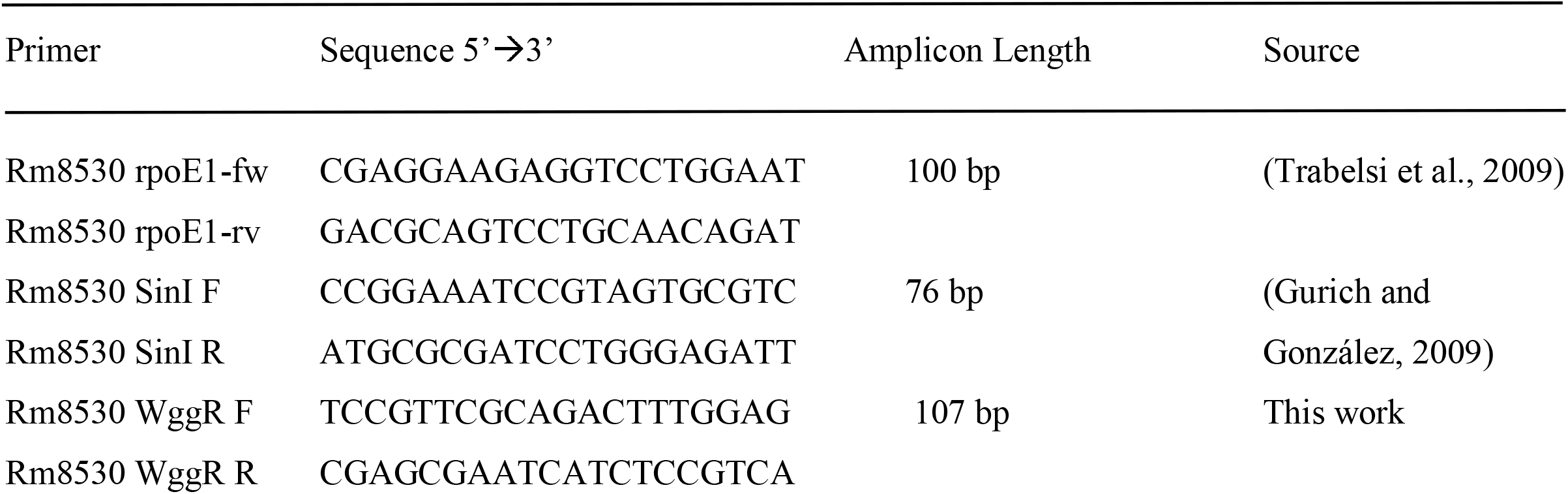
Primer sequences used in this study.

### Experimental Protocol for qRT-PCR

For qRT-PCR analysis, cells were ‘pre-conditioned’ on MGM media as described under ‘Biofilm Assays’ section. Cells were pelleted and re-suspended in fresh MGM plus the treatment. Co-inoculations were combined as Rm8530 OD_600_ = 0.8 and UD1022 OD_600_ = 0.2. Luteolin treatments contained a final concentration of 5 µM luteolin. Treatments were grown shaking at 30° C, and 1.5 mL samples were collected at time points of 12 and 24 hours, centrifuged, decanted and flash frozen in liquid nitrogen. RNA was isolated using NucleoSpin^®^ RNA from Macherey-Nagel (Düren, Germany). cDNA was generated with 500 ng RNA using High Capacity cDNA Reverse Transcription Kit from Applied Biosystems (www.appliedbiosystems.com) and qPCR was performed using PerfeCTa^®^ SYBR^®^ Green SuperMix, ROX, Quanta Biosciences (Gaithersburg, MD), and run on Eppendorf Mastercycler^®^ ep *realplex*^2^ (www.eppendorf.com). Experiments were repeated three times.

### Expression Analysis of qRT-PCR

The relative change in gene expression was calculated with the 2 ^-Δ Ct^ method as described in (Schmittgen and Livak, 2008), which calculates the expression of the gene of interest relative to the internal control in the treated sample compared with the untreated control. The internal control gene for Rm8530 is *rpo*E1. Genes were considered to be differentially expressed if the fold change in expression was ≥ 2 or ≤ -2.

### AHL Biosensor Assays for Quorum Quenching Analysis

Preparation of the AHL biosensor *Agrobacterium tumefaciens* KYC55 was as described in (Joelsson and Zhu, 2005) with modifications. KYC55 pre-induced cells were inoculated 1:1,000 into MGM medium for X-Gal soft agar 6-well plates. Pre-induced KYC55 cells were made as described in (Joelsson and Zhu, 2005). Soft agar plates were treated the same day they were poured. UD1022 was inoculated from a fresh plate streaked from glycerol stock into TYC and grown shaking 30° C for 5 hours to OD_600_ = 1.5, then sub-cultured 1:100 to MGM media and grown shaking 30° C for 20 hours to OD_600_ = 0.5. Treatments were made using these cultures mixed into sterile micro-centrifuge tubes with standard C8-AHL and 3-oxo-C16-AHL to a final concentration of 10 µM in volume of 200 µL. Controls contained standard AHL only. Treatments were incubated shaking 30° C for 24 hours.

Samples were then centrifuged at 16.1 *x 10,000 g* for 10 minutes at 4° C. Supernatants were transferred to new sterile tubes and sterilized open in biosafety cabinet under UV light for 30 minutes. 2 µL of treatments were applied to KYC55 X-Gal soft agar 6-well plates and allowed to dry. Plates were sealed with Parafilm^®^ and incubated agar side up for 24 hours at 30° C. Two treatment replicates were included on 2 separate 6-well plates. AHL biosensor assay was repeated twice.

### Sequence homology and alignment

The FASTA protein sequence YtnP protein in *Bacillus subtilis* subsp. *subtilis* str. 168 (sequence NP_390867.1) were queried using tblastn search translated nucleotide databases using a protein query against *B. subtillis* UD1022 nucleotide reference sequence (NZ_CP011534.1). The protein sequence for UD10222 YtnP has 281 amino acids and has a molecular weight of 31.8 kDa. The alignment of UD1022 YtnP, AiiA, and other MBL sequences was performed in MEGA (Tamura et al., 2011) by using the software MUSCLE (Edgar, 2004).

### Construction of *ytnP* mutant

The *ytnP* gene disruption *Bacillus subtilis* subsp. *subtilis* trpC2 *ytnP::erm* (Koo et al., 2017) was obtained from the Bacillus Genetic Stock Center and transferred into *B. subtilis* UD1022 by SPP1 phage transduction (Yasbin and Young, 1974).

### YtnP protein expression and purification

The *B. subtilis* UD1022 *ytnP* specific sequence was submitted to University of North Carolina School of Medicine Center for Structural Biology (NIH grant P30CA016086) for protein expression and purification. Workers sent the sequence to GenScript for gene synthesis and subcloning into a pET expression vector that contains an N-terminal His tag followed by a TEV site for tag removal during purification (pET-28a(+)-TEV. The *ytnP*::*E. coli* construct expression was done at UNC using their autoinduction expression system. Purification was performed using a Ni-affinity step, TEV protease tag removal, subtractive Ni-affinity step to separate out the tag and size-exclusion chromatography to remove potential protein contaminants.

## Results

### Co-inoculation of UD1022 and Rm8530 do not synergistically promote plant growth

*M. truncatula* plants were co-inoculated with *B. subtilis* UD1022 and *S. meliloti* Rm8530 6 days after germination and analyzed 7 weeks after inoculation for biomass and nodulation. Though the co-inoculation of Rm8530 and UD1022 resulted in no statistical difference in shoot biomass (**Figure 1A**, *p* = 0.06), there was a slight decrease in observable shoot growth (**Figure 1C**). There was no statistical difference in nodule numbers (**Figure 1B**, *p* = 0.59) between co-inoculated plants from those inoculated with Rm8530 alone. These results indicate that the addition of the PGPR UD1022 to the symbiotic strain Rm8530 did not increase plant health, contrary to results from similar studies (Fox et al., 2011; Morel et al., 2015). We posited that the lack of growth promotion by the co-inoculation may be due to antagonistic activity of UD1022 against Rm8530. A standard cross-streak compatibility assay on solid media determined no direct growth inhibitory effect between the two bacteria (**Supplementary Figure S1**).

**Figure 1.**
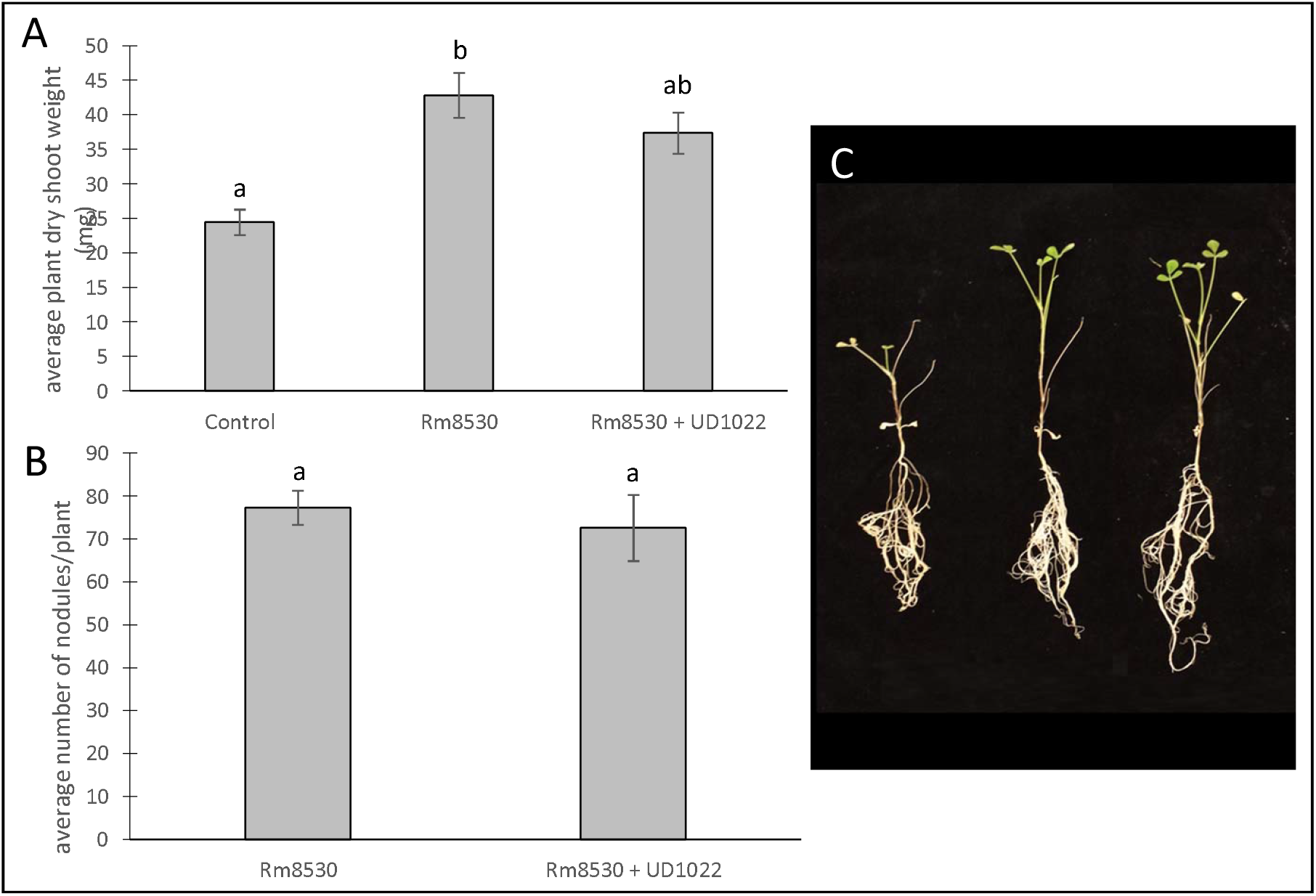
Co-inoculated plant growth and nodulation. **(A)** Average plant dry shoot weight of Rm8530 treated control plants and co-inoculated Rm8530 & UD1022 plants did not differ statistically (*p*-value of 0.06). **(B)** There was no statistical difference between average counts of nodules between treatments (*p*-value of 0.59). **(C)** Overall plant growth of both treatments was greater than control (first plant), but no differences were observed between Rm8530 treatment (second plant) and Rm8530 & UD1022 co-inoculation (third plant).

### UD1022 interacts indirectly with Rm8530 by interfering with Rm8530 biofilm and quorum sensing

UD1022 had no observable direct effects on Rm8530 growth; consequently, treatments of UD1022 culture filtrate supernatant (CFS) were tested for indirect influence on the Rm8530 functional phenotype of biofilm production. Biofilm formation by the symbiotic Rm8530 strain is required for efficient nodulation (González et al., 1996) and was evaluated by the semi-quantitative O’Toole assays (O’Toole et al., 2000) in treatments with UD1022 culture filtrate supernatant (CFS). Biofilm of Rm8530 cultured with 5% by volume of UD1022 CFS was significantly reduced from that of control (**Figure 2**, *p* <0.0001). Growth of Rm8530 with heat treated CFS treatment resulted in restoration of control quantities of biofilm (Figure 2, *p* = 0.86), suggesting the active factor of UD1022 CFS may be a protein.

**Figure 2.**
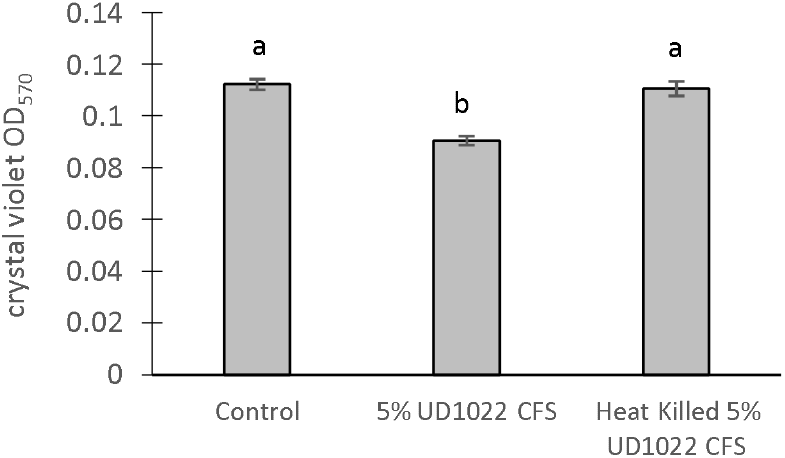
Rm8530 biofilm formation assay. Treatment with 5% UD1022 CFS significantly reduced the formation of biofilm by Rm8530 (*p*-value of <0.0001). Treatment with ‘heat killed’ UD1022 CFS showed no significant difference compared to the control (*p*-value of 0.86).

### UD1022 affects Rm8530 QS-controlled biofilm gene expression

The gene expression pathway for biofilm formation in *S. meliloti* is well characterized and tightly regulated through the ExpR/SinI QS system (Pellock et al., 2002; Marketon et al., 2003). The expression of two key Rm8530 QS genes were measured in response to co-culture with UD1022 CFS and in co-culture with live UD1022 cells using quantitative RT-PCR (qRT-PCR). The *sinI* gene encodes the QS autoinducer synthase SinI, which synthesizes unique long-chain N-acyl-homoserine lactone (AHL) signal molecules (Marketon et al., 2002). WggR is a transcriptional regulator that activates the downstream *wge* operons responsible for the biosynthesis and polymerization of EPS II low molecular weight galactoglucans, the symbiotically active exopolysaccharide component of *S. meliloti* biofilm (Gao et al., 2012).

Rm8530 *sinI* relative gene expression increased by 4-fold and *wggR* relative gene expression decreased by nearly 3-fold in treatments grown with UD1022 (**Figure 3**). These UD1022 live-cell co-culture qRT-PCR results reflected the same trend of expression as observed in the GFP gene expression reporter assays treated with UD1022 CFS (**Figure 4**): upregulation of *sinI* and downregulation of *wggR*. Treatments with the *M. truncatula* specific flavonoid luteolin (Peters et al., 1986) were included in the qRT-PCR expression analysis to evaluate possible plant host role in the interaction of the bacteria. Luteolin induces *nod* gene expression in *S. meliloti*, an important initial signaling mechanism to initiate legume – bacteria symbiosis. Rm8530 QS gene expression, as expected was not directly affected by the presence of luteolin alone. However, the presence of luteolin in Rm8530 – UD1022 co-culture significantly enhanced the gene expression changes observed in bacteria co-cultures. The increase in *sinI* relative expression doubled to nearly 8-fold and *wggR* decreased expression was extended to 3.4-fold (**Figure 3**). This could indicate that, in the rhizosphere, plant signaling factors such as flavonoids could exacerbate the PGPR interactions causing the changes in QS gene expression.

**Figure 3.**
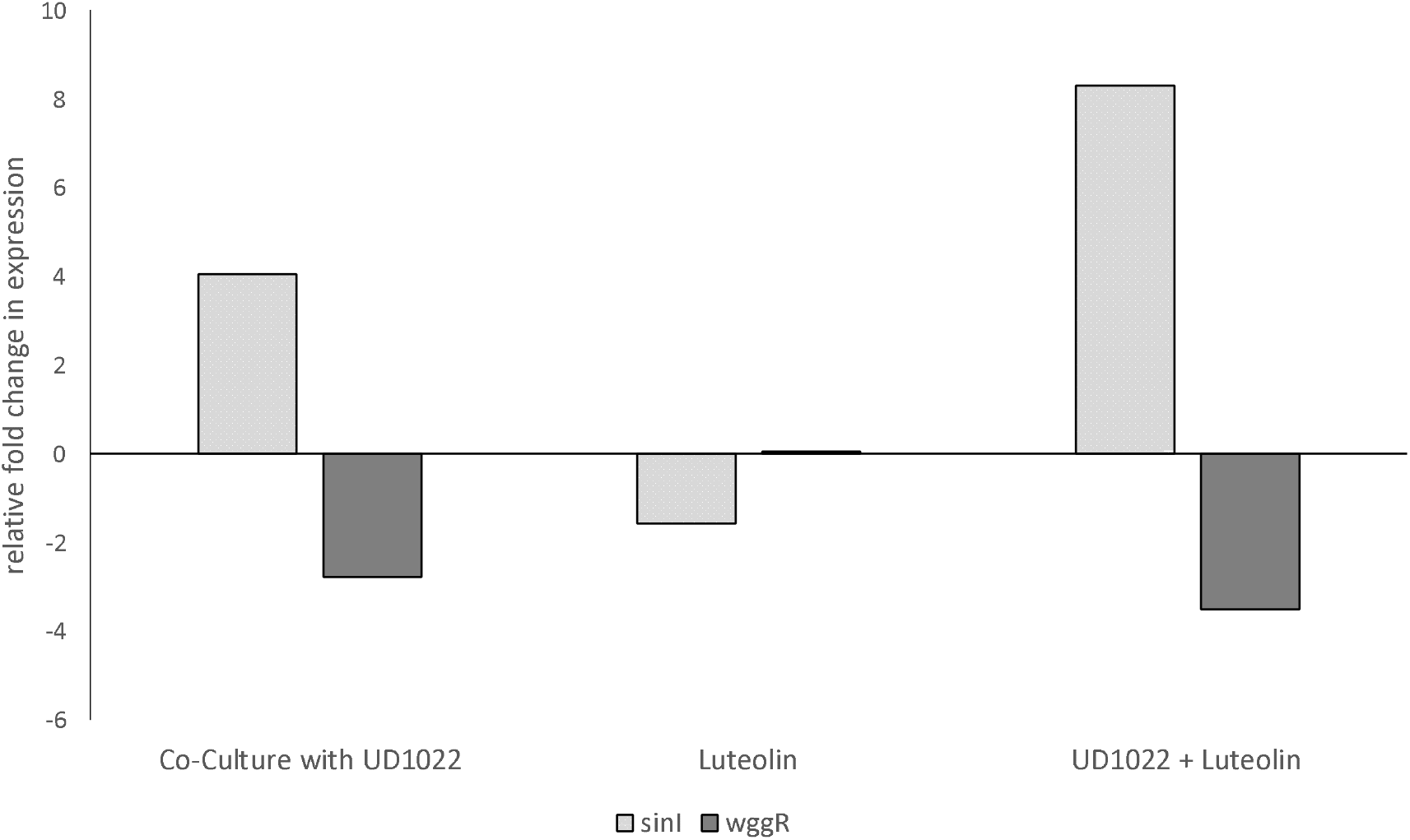
Relative fold changes in expression of Rm8530 *sinI* and *wggR* in co-culture with UD1022. Co-culture with UD1022 increased the relative expression of Rm8530 *sinI* by 4-fold. The presence of luteolin (which represents the condition of Rm8530 upregulating Nod genes) doubled the effect of UD1022 on Rm8530 *sinI*, increasing expression to 8-fold. Luteolin alone did not significantly change Rm8530 *sinI* expression. Co-culture with UD1022 decreased the relative expression of Rm8530 *wggR* by almost 3-fold. The presence of luteolin slightly enhanced the effect of UD1022 on Rm8530 *wggR*, increasing to 3.5-fold. Luteolin alone did not significantly change Rm8530 *wggR* expression.

**Figure 4.**
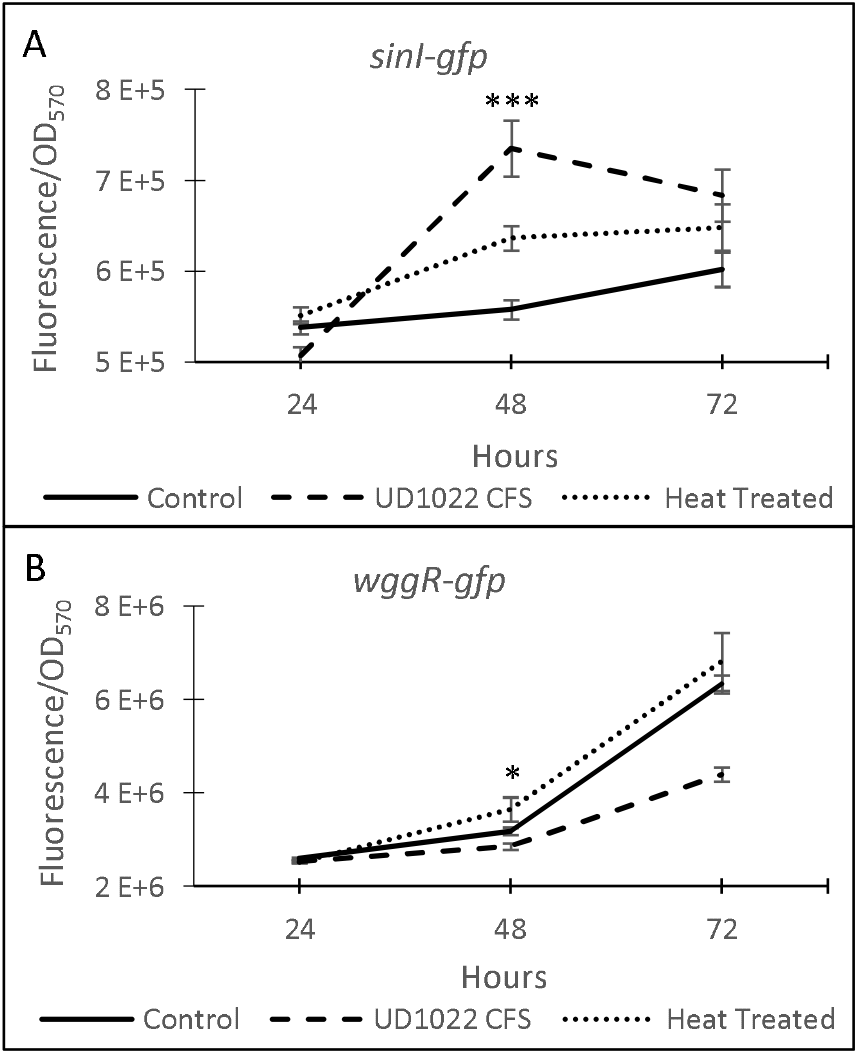
Expression of Rm8530 quorum sensing genes. **(A)** Average GFP activity (fluorescence/OD_570_) of the *sinI-gfp* fusion reporter. The differences between treatments at 48 hours is significant at *p*-value of <0.0001. **(B)** Average GFP activity of *wggR-gfp* fusion reporter. The differences between the treatments and the control at 48 hours is significant at *p*-value of 0.04, and 72 hours is significant with *p*-value of <0.0001. Averages for both assays are from 29 technical replicates and the bars at each time point present standard error. * indicates *p* < 0.05 and *** *p* < 0.0001 in Kruskal-Wallis test.

The regulatory network of *S. meliloti* ExpR/SinI is intricately controlled through AHL acyl chain length, acyl chain substitutions, and concentration of AHL molecules (Bartels et al., 2007; Calatrava-Morales et al., 2018). Genes expression for *sinI* synthase is positively regulated by low concentrations of AHLs (1-40 nM) and negatively regulated by high concentrations of AHLs (>40 nM), allowing the ExpR transcriptional regulator of *sinI* to be sensitive to the AHL substrate it is responsible for producing (Baumgardt et al., 2014). The expression of *wggR* is oppositely regulated, requiring upwards of 150 nM C_16:1_ -AHL for increased *wggR* GFP expression reporter activity (Gao et al., 2012). Lower concentration of AHLs in UD1022 treatments would support the patterns of increased *sinI* and decreased *wggR* expression (schematic in **Figure 5**).

**Figure 5.**
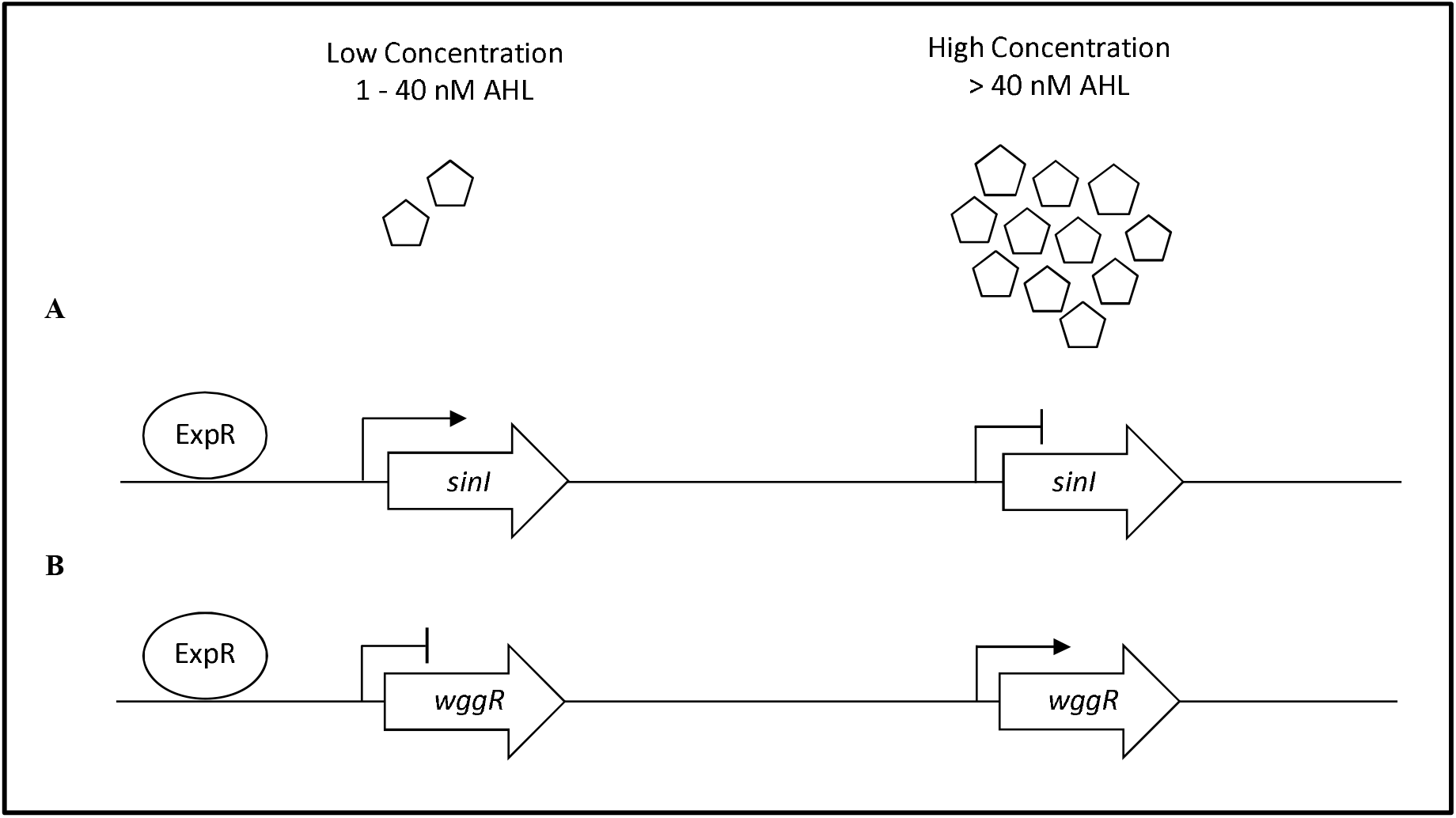
Generalized *sinI*/ExpR quorum sensing transcription model for *sinI* and *wggR*. Expression of genes are controlled on multiple levels including length of AHL chain and AHL concentration. **(A)** Low concentrations of AHL (1-40 nM) upregulate *sinI* expression in a positive feedback regulation. **(B)** Low concentrations of AHLs downregulate *wggR* expression. Adapted from (Baumgardt et al, 2014).

### UD1022 is positive for quorum quenching (QQ) activity against Rm8530

The Rm8530 QS gene expression response patterns coupled with the restoration of WT Rm8530 biofilm formation in UD1022 heat treated CFS treatments suggest that UD1022 may be affecting Rm8530 QS through enzymatic activity of a protein. Interference of QS through interspecific enzymes is termed quorum quenching (QQ) and can be enacted at several levels of QS regulation, including targeting signal biosynthesis, signal receptors and direct cleavage of QS signal molecules, including AHLs (Fetzner, 2015). QQ enzymes have been characterized in many soil bacteria including *Agrobacterium* and *Bacillus* genera (Chan et al., 2016).

QQ activity of UD1022 was assessed using the bioreporter strain *Agrobacterium tumefaciens* KYC55 to detect a wide range of AHLs and reports through β-galactosidase activity (Zhu et al., 2003). UD1022 cultures were incubated with 10 µM purified *N*-octanoyl-L-homoserine lactone (C8-AHL) or N-3-oxo-hexadecanoyl-L-homoserine lactone (3-oxo-C16-AHL) (Caymen Chemicals) for 24 hours on KYC55 X-Gal plates. Treatments of UD122 with 3-oxo-C16-AHL showed significant reduction in detectable AHL signal (**Figure 6B**.), while C8-AHL showed no discernable difference from the control (**Supplementary Figure S2B**). Thus, UD1022 displays quorum quenching activity which appears to be geared toward long-chain AHLs.

**Figure 6.**
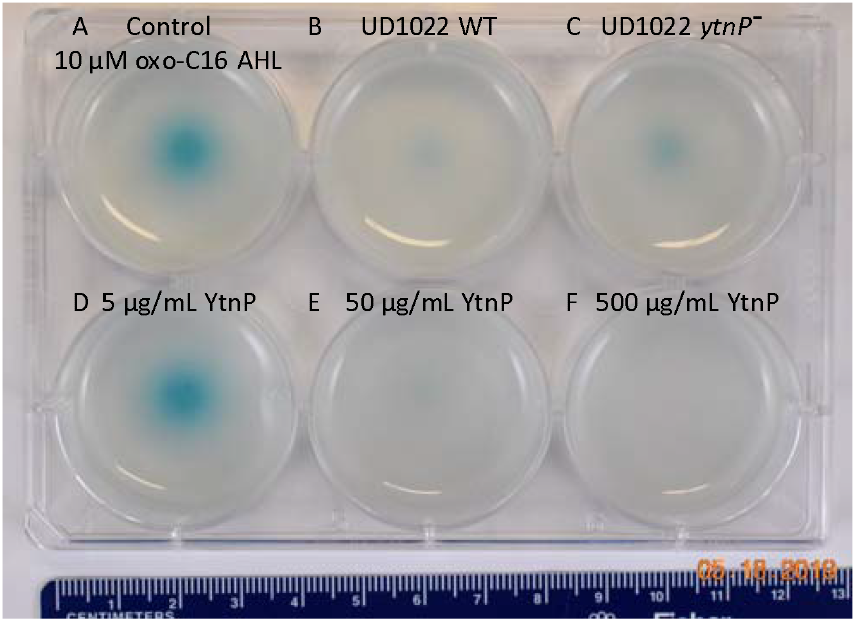
UD1022 Quorum quenching biosensor assay plate. The biosensor KYC55-X-gal soft agar plate treated with UD1022-AHL co-cultures. From top left across: **(A)** control treatments of standard AHLs with no UD1022. **(B)** QQ activity of UD1022 culture with 3-oxo-C16-AHL **(C)** UD1022 *ytnP*^—^ mutant cultured with AHL **(D)** 5 µg/mL pure UD1022 YtnP protein incubated with AHL, **(E)** 50 µg/mL YtnP protein, **(F)** 500 µg/mL YtnP protein. *Brightness of image increased by 20%, which did not increase pigment intensity or saturation.

### UD1022 QQ through the lactonse YtnP protein

Several classes of bacterial enzymes are known for QQ through inactivating AHLs, including lactonases and acylases (Chan et al., 2016). A search of the literature for lactonases specifically identified in *B. subtilis* species yielded the putative lactonase YtnP protein in *B. subtilis* NCIB3610 (Schneider et al., 2012). The alignment of YtnP protein sequence with UD1022 returned a 98.44% identity. Using MUSCLE (Edgar, 2004) alignments of the reference protein *B. subtilis* strain 168 YtnP (NP_390867.1) and UD1022 YtnP sequences revealed the hallmark metallohydrolase HXHXDH and HXXGH metal binding motifs as well as a phosphorylated Ser36 residue. To determine if the UD1022 YtnP lactonase protein contributes to the QQ patterns observed, we introduced an *ytnP* deletion cassette in UD1022 (‘UD1022 *ytnP*^—^’). In AHL co-incubation assays, UD1022 *ytnP*^—^ treatments with 3-oxo-C16-AHL showed that AHL degradation was less extensive as that of UD1022 WT (**Figure 6C**). It is likely that there are additional QQ active proteins produced by UD1022. Indeed, up to six other probable MBL-like fold sequences having the HXHXDH motif have been identified in UD1022 (data not shown).

The UD1022 specific *ytnP* sequence was submitted to the University of North Carolina School of Medicine Center for Structural Biology (NIH grant P30CA016086) for protein expression and purification. Purified YtnP protein was applied at 3 different concentrations to 10 µM concentrations of 3-oxo-C16-AHL and C8-AHL. The biosensor reporter showed no degradation of 3-oxo-C16-AHL with treatment of 5 µg/mL YtnP (**Figure 6D**). Long chain AHL degradation comparable to UD1022 WT live cell treatments was observed with 50 µg/mL YtnP incubation (**Figure 6E**). Treatments of 500 µg/mL YtnP completely abolished detectable levels of 3-oxo C16-AHL (**Figure 6F**). Incubation of UD1022 YtnP with C8-AHL, interestingly, resulted in degradation of the short-chain AHL at 50 µg/ml and 500 µg/mL (**Supplementary Figure S2E** and **S2F**, respectively). This demonstrates unequivocally that UD1022 QQ activity is carried out through the YtnP lactonase protein.

## Discussion

Understanding interactions of PGPR in consortia is critical for predicting rhizo-microbiome function in the environment and in agroecosystems. This is especially relevant as biological-based crop solutions become more widely marketed and adopted. Several examples of PGPR co-inoculations using *S. meliloti* resulting in significant improvements of *Medicago* spp. plant growth have been reported. Co-inoculation of *Delftia* spp. JD2, a diazotrophic, IAA producing PGPR, with *S. meliloti* U143 onto *M. sativa* increased nodulation (Morel et al., 2011), and increased shoot and root dry weights by 13% and 34%, respectively (Morel et al., 2015). Fox et al. (2011) found duel inoculation of *S. meliloti* WSM419 and the PGPR *Pseudomonas fluroescens* WSM3457 onto *M. truncatula* enhanced nodule initiation rates, resulting in increased number of crown nodules and more overall N accumulation. *S. meliloti* B399, a commercial alfalfa inoculant closely related to strain Sm1021, co-inoculated with *Pseudomonas* spp. FM7d nearly doubled shoot dry weight and increased nodule number on *M. sativa* L. cv Bárbara SP (INTA Manfredi). Though co-inoculation of B399 with *Bacillus* spp. M7c had significantly higher shoot dry weight, it did not increase nodule number (Guiñazú et al., 2010).

However, not every instance of PGPR co-inoculation with *S. meliloti* has been reported to be beneficial. The duel inoculation of the PGPR *B. simplex* 30N-5 with *S. meliloti* 1021 onto *M. truncatula* resulted in no significant difference in shoot height, plant dry weight or nodule number over that of *S. meliloti* 1021 control (Maymon et al., 2015). This contrasted with their previous work which showed beneficial growth effects of *B. simplex* 30N-5 when co-inoculated with *Rhizobium leguminosarum* bv. *viciae* 128C53 onto pea (*Pisum sativum*) (Schwartz et al., 2013). Our study using the *expR*+ *S. meliloti* strain Rm8530 co-inoculated with *B. subtilis* UD1022 also resulted in no significant enhancement of plant growth or nodule number. While other work has yet to query the mechanisms of bacterial interaction which may account for the non-synergistic plant effects of these rhizobia-PGPR co-inoculations, this work reveals a potential, indirect mechanism of bacterial interaction.

Biofilm formation is important in soil and root associated bacteria for motility and exchange of signals and metabolites (Angus and Hirsch, 2013; Bogino et al., 2013; Amaya-Gómez et al., 2015). *S. meliloti* biofilms have been shown to play a critical role in motility toward and initiation of nodulation with the *Medicago* spp. plant root (González et al., 1996; Pellock et al., 2000; Hoang et al., 2008). Here we used Rm8530 biofilm formation as functional reporter for negative activity by UD1022 and found clear evidence of UD1022 inhibition of Rm8530 biofilm formation. *S. meliloti* biofilm formation is dependent on intact ExpR/SinI QS system, which is well described for both strains Rm1021 and its ExpR+ relative Rm8530. Importantly, the Rm8530 QS system has been shown to regulate a key symbiotically active component of their biofilms, the low molecular weight galactoglucans referred to as EPS II (Rinaudi and Gonzalez, 2009).

Based on the negative effect of UD1022 on Rm8530 biofilm, we hypothesized that UD1022 may be interfering with the QS controlled molecular regulation of biofilm production. The Rm8530 QS genes *sinI* and *wggR* were selected to test the effect of UD1022 on the QS pathway, including upstream QS signal molecule synthesis (*sinI*) and downstream EPS II polymerization (*wggR*). Using UD1022 CFS treatments on Rm8530-*gfp* expression reporters and subsequent validation with qRT-PCR of live-cell co-cultures, we found UD1022 significantly activated *sinI* transcription and reduced *wggR* transcription. McIntosh et al. (2009) described that *sinI*-promoter activation occurs at nearly 10-fold lower levels of AHLs than required for its downregulation. WggR activation requires the presence of the transcriptional regulator ExpR and the SinI specific AHLs C_16:1_-AHL and oxo-C_16:1_-AHL (McIntosh et al., 2009; Gao et al., 2012). The *wggR*-*gpf* reporter in Rm8530 *sinI* background was more sensitive to C_16:1_-AHL than 3-oxo-C_16:1_-AHL. Expression of *wggR* increased in a dose dependent manner with close to WT levels at 40-1500 nM C_16:1_-AHL and 200-1500 nM oxo-C_16:1_-AHL (Gao et al., 2012). Consequently, UD1022 appeared to stimulate an expression pattern of the QS genes similar to their response to low AHL signal molecule concentration conditions.

QQ activities could be promising as a prospective tool to improve plant health and bypass antibiotic resistance in the development of biological products combating plant pathogens (Grandclément et al., 2016; Rodríguez et al., 2020). Many screening techniques have been utilized to identify QQ microbial isolates for this purpose (Tang et al., 2013; Last et al., 2016; Stein and Schikora, 2018). To better understand the capacity of UD1022 for QQ, we used the bioreporter strain *Agrobacterium tumefaciens* KYC55 in soft-agar to detect both short and long-chain AHLs. When co-cultures of UD1022 were incubated with 10 µM purified 3-oxo-C16-AHL for 24 hours and applied to the bioreporter, expression of KYC55 β-galactosidase was greatly diminished. This reduction of detectable long chain AHL demonstrates that UD1022 is capable of QQ activity. Response of the bioreporter to co-cultures of UD1022 with C8-HSL was no different from control treatments.

Many modes of QQ by bacteria have been identified with lactonase hydrolytic enzymes being highly described in *Bacillus* spp. (Kumar et al., 2015). The *B. subtilis* NCIB3610 putative lactonase YtnP protein sequence had high similarity to that found in UD1022 (Schneider et al., 2012). The YtnP lactonase is a metallolactamase and was found to target γ-butyrolactone of *Streptomyces griseus*. The UD1022 YtnP protein possesses the same hallmark metallohydrolase features of the NCIB3610 YtnP, including the HXHXDH motif, indicating that UD1022 YtnP is also a likely a QQ lactonase. The UD1022 *ytnP*^—^ mutant was employed in the AHL biosensor assay to test the role of the specific YtnP QQ activity. Rather than fully abolishing QQ, the partial degradation activity remaining may be due to redundant or multiple lactonase-like genes which continue to be expressed in the single *ytnP* mutant. The purified YtnP protein incubated with long and short chain AHLs showed clear and efficient QQ activity. Exogenous application of pure YtnP degraded C8-AHL, which was not observed in UD1022 live-cell assays. The substrate specificity of the YtnP lactonase protein may be more reliant on cognate enzyme:substrate ratios rather than primarily on acyl-chain length of the AHL. Lactonases characterized to date are described as having broad activity against a range of AHL acyl chain lengths and substitutions, though with variable active site affinities (Bergonzi et al., 2018). No enzyme kinetics or substrate specificity analysis were performed for this work, but the perceived biological inactivity on short chain AHL may be attributed to a biological interaction of substrate concentration or may be an anomaly due to KYC55 variation in response sensitivity to different AHLs (Zhu et al., 2003).

QQ in the rhizosphere likely plays a large part in PGPR interactions and, consequently, in plant health outcomes. The presence of AHL molecules in the rhizosphere have been shown to directly elicit functional and beneficial responses from both non-legumes and legumes (Hartmann et al., 2014; Schikora et al., 2016; Hartmann and Rothballer, 2017). A recent study expanded on previous results showing the direct application of long-chain AHLs increased the number of nodules on *M. truncatula* inoculated with *S. meliloti* 1021 by showing this phenomenon occurred only when seeds were pre-treated with antibiotics (Veliz-Vallejos et al., 2020). This study illustrates the contribution of plant bacterial consortia toward altering beneficial outcomes involving QS signals. Several studies have employed the use of bacterial QQ lactonases to demonstrate the direct and indirect beneficial activities of AHLs and QS on plants. To identify novel QS controlled proteins, an *S. meliloti* 1021 construct expressing the QQ lactonase AiiA was found to be significantly deficient in forming nodule initials within the first 12 hours after inoculation (Gao et al., 2007). Zarkani et al. (2013) showed that *S. meliloti* producing 3-oxo-C14-AHL increased *Arabidopsis thaliana* resistance to *Pseudomonas syringe* pv *tomato* DC3000, while mutants heterologously expressing the *Agrobacterium tumefaciens* AttM lactonase did not.

As discussed in previous sections above, the Rm8530 QS system is required to produce symbiotically active EPS II biofilms (Pellock et al., 2000; Pellock et al., 2002; Hoang et al., 2004; Gurich and González, 2009). QS is also important in controlling bacterial cell population density, motility toward the plant root and switching expression pathways from motility to nodulation (Bahlawane et al., 2008; Gao et al., 2012; Calatrava-Morales et al., 2018). The timing and coordination of these activities is intricately controlled through ExpR/SinI QS and disruptions or interference through QQ has the potential to affect the efficiency and competency of these pathways. The lack of synergistic effects between Rm8530 and UD1022 may be explained, in part, through the QQ activity of UD1022 YtnP lactonase reducing Rm8530 AHL signal molecule concentration, leading to reduced expression of EPS II biosynthesis genes including *wggR*, and ultimately resulting in inhibition of efficient nodule initiation on *M. truncatula* roots (**Figure 7**).

**Figure 7.**
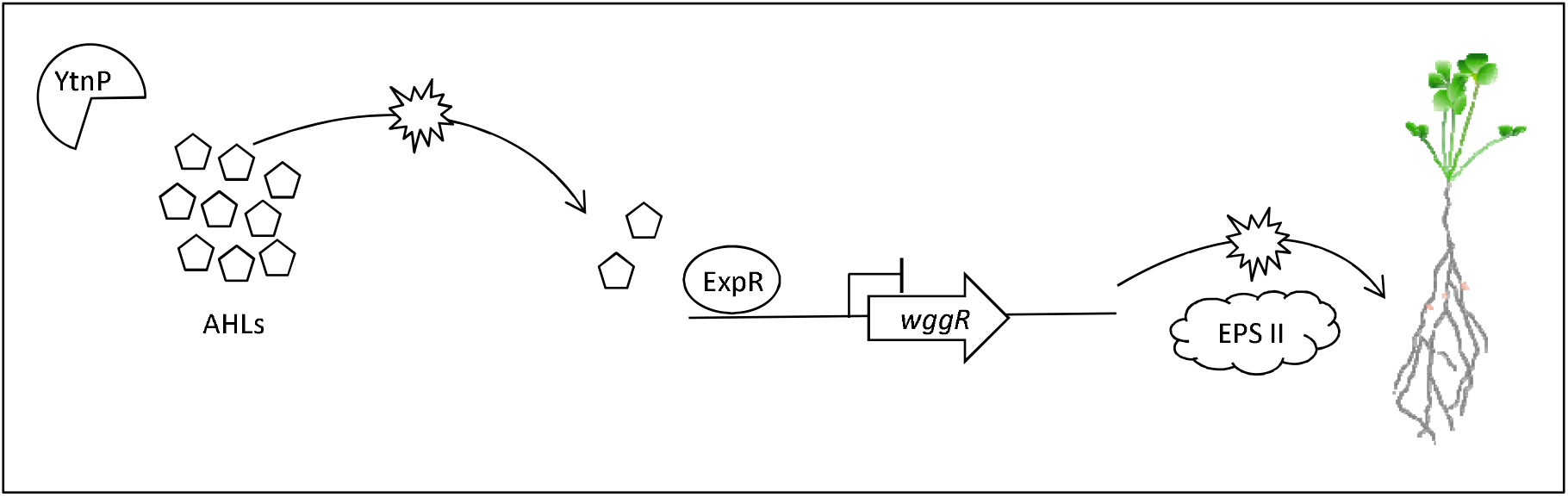
Model of proposed molecular QS and QQ interactions between *M. truncatula* PGPRs. *B. subtilis* UD1022 produces the lactonase YtnP which cleaves *S. meliloti* Rm8530 AHLs. Through quorum quenching, UD1022 YtnP reduces AHL concentrations, inhibiting the upregulation of symbiotically active EPS II genes. This may result in lower nodulation efficiency of Rm8530 in the presence of the PGPR UD1022.

## Conclusion

We show in our tri-trophic legume – symbiont – PGPR model system that the PGPR *B. subtilis* UD1022 does not synergistically increase *M. truncatula* plant growth or nodulation by the legume symbiont *S. meliloti* Rm8530. Though there is no direct growth inhibitory effect between the bacterial strains, indirect interactions contribute to the disruption of plant associative activities by the symbiont. UD1022 affects Rm8530 QS controlled biofilm formation through interference with the QS biosynthesis pathway. Further, UD1022 expresses the QQ lactonase YtnP, which cleaves the specific AHLs required to produce symbiotically active EPS II of Rm8530 biofilms. UD1022 likely delays or fails to promote Rm8530 nodulation through the QQ activity of lactonase YtnP and can inhibit synergistic plant growth promotion.

## Data Availability Statement

The raw data supporting the conclusions of this article will be made available by the authors, without undue reservation.

## Author Contributions

All authors designed the experiments, edited, and contributed to the final manuscript, and approved for submission. AR completed lab work pertaining to microscopy and genetic expression. AR and HPB conceptualized the idea, PBB designed the experiments for the *Bacillus* mutants.

## Acknowledgments

We thank Dr. Juan E. González for providing *S. meliloti* strain Rm8530, Dr. Max Teplitski for providing *S. meliloti* Rm8530 *sinI*-*gfp* and *wggR*-*gfp* strains, Dr. Stephen Winans for providing *Agrobacterium tumefaciens* KYC55 (pJZ410)(pJZ384)(pJZ372), and the University of North Carolina School of Medicine Center for Structural Biology for UD1022 *ytnP* protein expression. The results, in part, are based on work reported in the master’s thesis of Amanda Rosier, which is freely available through the University of Delaware Library Institutional Repository (http://udspace.udel.edu/bitstream/handle/19716/21138/2016_RosierAmanda_MS.pdf).

