## Supplementary Figures for "Quorum quenching activity of the PGPR *Bacillus subtilis* UD1022 alters nodulation efficiency of *Sinorhizobium meliloti* on *Medicago truncatula*"

Supplementary Material

**Supplementary Figures**

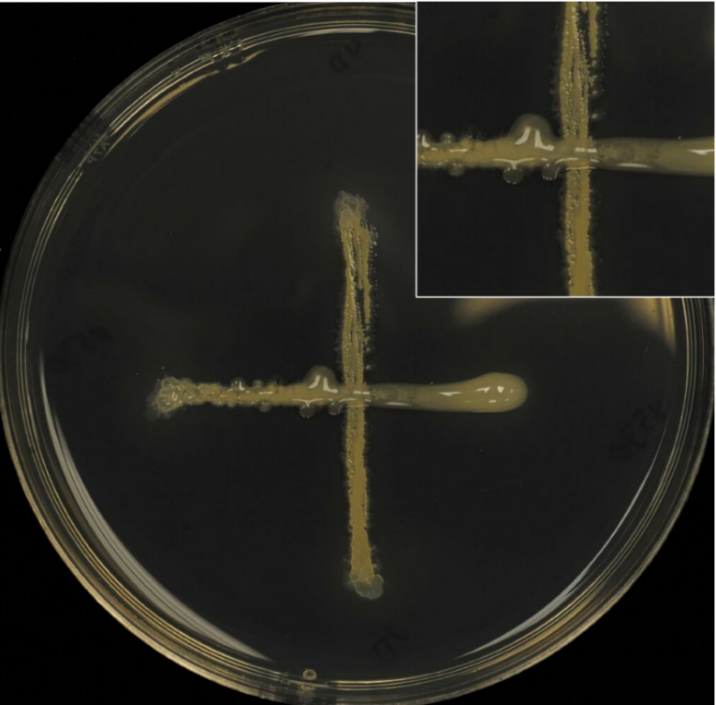

**Supplementary Figure 1.** Cross-streak assay. The bacteria *S. meliloti* strain Rm8530 (horizontal) and *B. subtilis* strain UD1022 (vertical), were streaked together to test for growth inhibition. The two bacteria demonstrated no growth inhibition on solid or in liquid media (data not shown).

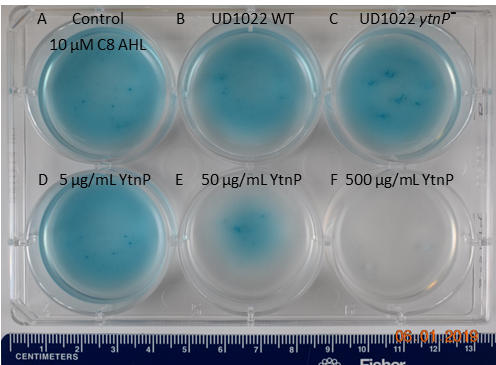

**Supplementary Figure 2.** UD1022 Quorum quenching biosensor assay plate. The biosensor KYC55-X-gal soft agar plate treated with UD1022-AHL co-cultures. From top left across: **(A)** control treatments of standard AHLs with no UD1022. **(B)** QQ activity of UD1022 culture with C8-HSL **(C)** UD1022 *ytnP*¯ mutant cultured with AHL **(D)** 5 µg/mL pure UD1022 YtnP protein incubated with AHL, **(E)** 50 µg/mL YtnP protein, **(F)** 500 µg/mL YtnP protein.
